# Molecular and Functional Asymmetry in Cckar-expressing Vagal Sensory Neurons

**DOI:** 10.1101/2024.11.01.621506

**Authors:** Hailey F. Welch, Ishwarya Sankaranarayanan, Veronica M. Hong, Haniya Qavi, Khadijah Mazhar, Benedict J. Kolber, Theodore J. Price, Catherine A. Thorn

## Abstract

The vagus nerves are important carriers of sensory information from the viscera to the brain. Emerging evidence suggests that sensory signaling through the right, but not the left, vagus nerve evokes striatal dopamine release and reinforces appetitive behaviors. However, the extent to which differential gene expression within vagal sensory neurons may underlie this asymmetric reward-related signaling is unknown. Here, we use single-nucleus RNA sequencing, *in situ* hybridization, and calcium imaging to identify a single cluster of neurons, defined by co-expression of *Chrna3* (nicotinic acetylcholine receptor subunit 3) and *Cckar* (cholecystokinin 1 receptor), that is preferentially expressed in the right nodose ganglia of rats and mice. This neuronal population also expresses several genes implicated in digestive signaling, including *Glp1r* and *Sctr*, consistent with a gut-innervating subtype. Our results suggest that right-biased expression of this unique chemosensing vagal cell type may contribute to asymmetric encoding of interoceptive rewards by the vagus nerves.

## INTRODUCTION

The vagus nerves (cranial nerve X) transmit sensory information from the viscera, including cardiovascular, respiratory, and gastrointestinal (GI) organs, to the central nervous system to maintain homeostatic control over autonomic processes such as heart rate, respiration, and digestion ^1–3^. The sensory neurons of the vagus nerve are situated in the jugular and nodose ganglia (NG), located near the base of the skull. These neurons detect a wide range of visceral signals, including mechanical stretch and chemical stimuli, and relay this information to the brainstem to mediate appropriate autonomic and behavioral responses ^4–8^.

Several studies have demonstrated striking left-right asymmetries in sensorimotor vagal pathways, with the left and right vagus nerves exhibiting distinct patterns of innervation across visceral organs, including the heart, lungs, and gastrointestinal (GI) tract ^9–14^. These anatomical differences are thought to contribute to divergent functional outcomes following left vs. right vagal stimulation ^15–19^. Of particular interest is the emerging evidence for lateralized processing of visceral reward-related signals. Stimulation of the right vagus nerve, but not the left, is seen to reinforce behavior and activate midbrain dopaminergic reward nuclei ^18^, and gut-innervating vagal afferents likely play a key role in this lateralized reward signaling ^16^. These recent studies raise additional questions about the cellular origins of asymmetric reward encoding by the right versus left vagus nerve.

Genetic sequencing studies of the NG in mice have revealed incredible diversity among NG cell types ^6,20–23^. Transcription patterns of vagal neurons have been used to identify distinct subclasses of NG cells and their hypothesized sensory functions ^20,21,24^. This work has typically been performed on pooled samples from left and right NG, however, and cannot address the question of which NG genes or neuronal subtypes exhibit asymmetric left-right expression that might contribute to lateralized vagal signaling. Recently, a more targeted investigation of left vs. right NG expression of several genes associated with digestive signaling showed that, in mice, neurons expressing *Cckar* (cholecystokinin 1 receptor), *Glp1r* (glucagon-like peptide 1 receptor), and *Npy2r* (neuropeptide Y receptor Y2; peptide YY receptor 2) are more numerous in right NG compared to left ^12^. Differential left-right signaling through these receptors was not observed, however, leaving open the possibility of additional functionally relevant lateralized signaling pathways. In the current study, we use single-nucleus RNA sequencing (snRNA-seq) to identify genes of interest that are differentially expressed between the left and right NG that could contribute to right-biased vagus-mediated reward encoding. Because of their larger size compared to mice, rats are a more commonly used laboratory model to investigate translational vagal therapies such as vagus nerve stimulation (VNS). We thus performed comprehensive genetic profiling of left vs. right NG neurons in rats and compare our results to a publicly available mouse dataset ^25^. Computational results were functionally validated using RNA fluorescence *in situ* hybridization (RNA-FISH) and calcium imaging.

Our results identify a single cluster of neurons, marked by co-expression of *Cckar* and *Chrna3*, that is preferentially found in the right NG of both rats and mice. Our data further suggest that this right-biased neuronal subtype exhibits a genetic profile consistent with a gut-innervating population, including co-expression of *Glp1r* (glucagon like peptide 1 receptor) and *Sctr* (secretin receptor). We additionally find that population responses to cholecystokinin 1 receptor (CCK1R) activation likewise exhibit significant left-right asymmetry, with more neurons responding to CCK1R agonism in the right NG compared to left. These findings suggest that differential right > left expression of chemosensory vagal neurons expressing CCK1Rs could give rise to asymmetric transduction of visceral sensory signals by the vagus nerve. Combined, our study raises the possibility that right-biased digestive peptide signaling within this cell population, including at CCK1Rs, may contribute to the lateralized encoding of interoceptive rewards.

## RESULTS

### Diverse mechanosensitive and chemosensitive cell types identified in rat nodose ganglia

We performed snRNA-seq on 12 replicate groups of NG cell suspensions. Each replicate group was formed by pooling the right or left NG from six male or female rats (**Figure 1A**). In all, 36,741 nuclei were sequenced. Data were filtered according to standard quality control (QC) metrics, and data from all replicate groups were integrated prior to clustering (**Figure S1**) ^26,27^. Non-neuronal clusters (expressing *Ttn*, *Myh4*, *Mpz*, *Mbp*, *Apoe*, *Sparc*, *Emcn*, and/or *Tyrp1*) and jugular neuronal clusters (expressing *Pdrm12*) were identified and removed during an initial 4 rounds of cluster filtering (**Figure S2**). After QC and cluster filtering, the remaining 23,428 nuclei were reclustered, resulting in 19 clusters corresponding to putative NG neuron subtypes, which were included in subsequent analyses (**Figure 1B-C**).

**Figure 1.**
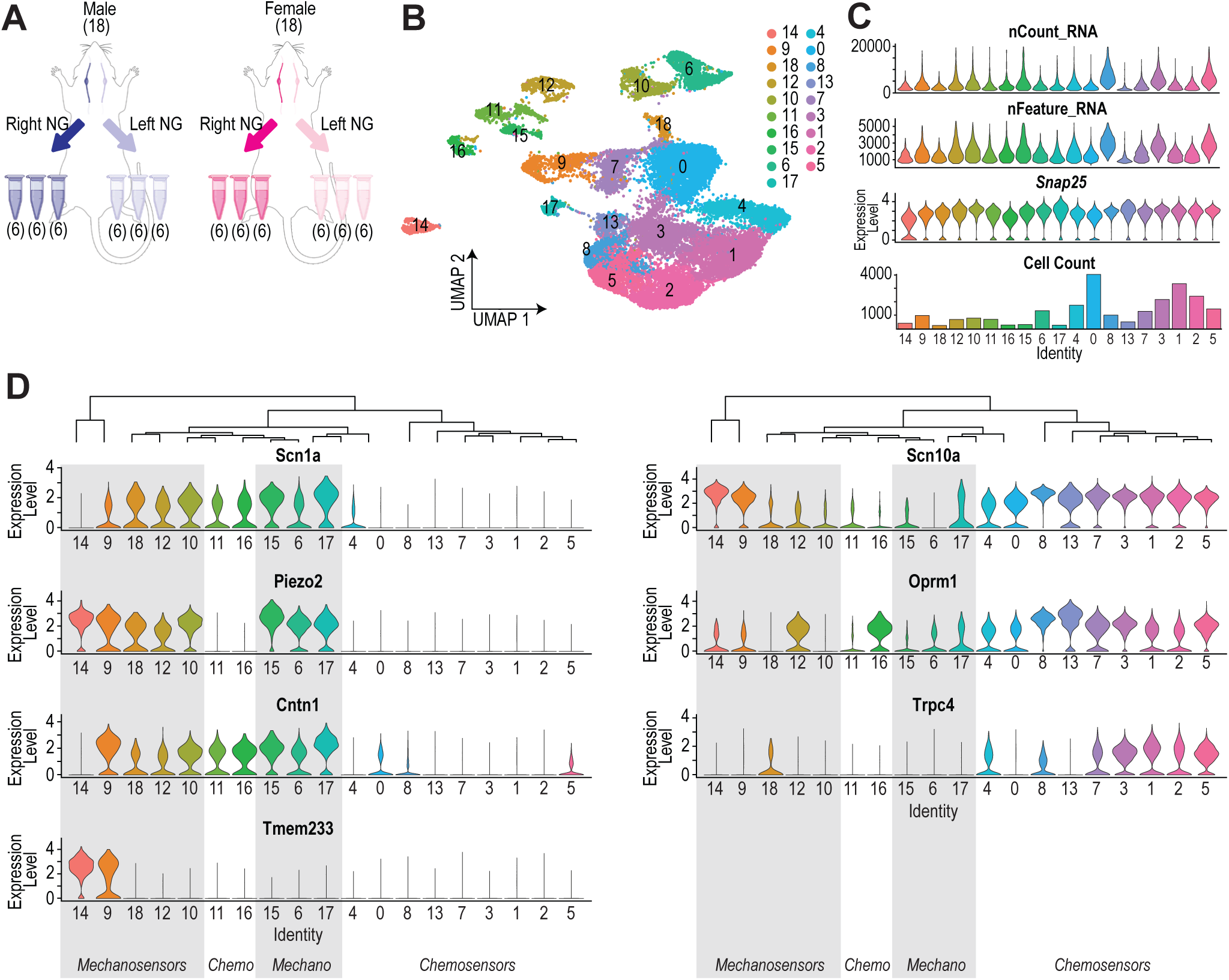
Putative mechanosensing and chemosensing clusters identified in left and right NG neurons in rats. A) Cartoon illustrating pooling of right and left NG from 18 male and 18 female rats into replicate groups. Created in BioRender. Thorn, C. (2024) BioRender.com/w17y601. B) After quality control and removal of non-neuronal and jugular ganglia clusters, 19 NG clusters were identified from integrated right and left samples (n = 23,428 nuclei). C) All NG clusters met quality control criteria and expressed high levels of the neuronal marker *Snap25*. D) Clusters can be divided into putative mechanosensing (gray boxes) versus chemosensing subtypes based on expression of *Piezo2.* Left: *Piezo2* expression overlaps with expression of *Scn1a and Cntn1* in most mechanosensory clusters; Right: In most chemosensory clusters, *Snc10a* expression overlaps with high levels of *Oprm1* and low levels of *Piezo2*.

Prior sequencing of mouse NG has identified *Piezo2* expressing (*Piezo2*+) putative mechanosensory cells and *Scn10a* expressing “nociceptor-like” cells in the NG ^21^. Our data, in rats, replicate these fundamental results. We observed minimal co-expression of two sodium channel genes, *Scn1a* (encoding Na_v_1.1) and *Scn10a* (encoding Na_v_1.8) (**Figure 1D**). Expression of *Scn1a* typically coincided with expression of *Piezo2* and *Cntn1* (**Figure 1D**, left), suggesting a myelinated and mechanosensory phenotype for the majority of these clusters. A smaller group of two *Piezo2+* putative mechanosensory clusters was also identified in our dataset, which preferentially expressed *Scn10a* over *Scn1a*, and co-expressed *Tmem233* (transmembrane protein 233). Most *Scn10a*-expressing clusters were not found to exhibit genes associated with mechanosensation, however. Rather, they co-expressed *Oprm1* (mu-1 opioid receptor) and many also co-expressed *Trpc4* (transient receptor potential cation channel subfamily C member 4), pointing to a chemosensory phenotype for the majority of *Scn10a*-expressing clusters (**Figure 1D**, right). Approximately 20% of the cells in our dataset (n = 5,006/23,428; 21.4%) were sorted into one of 8 *Piezo2*+, putative mechanosensory clusters, whereas the majority of NG cells (n = 18,422/23,428; 78.6%) were instead sorted into one of the 11 *Piezo2*-negative putative chemosensory clusters. Our computational results suggest that in rats, as has been shown previously in mice, there is little overlap of expression of *Scn1a* and *Scn10a* in NG neurons, and that expression of the *Scn1a* vs. *Scn10a* is highly (though not exclusively) correlated with putative mechanosensory vs. chemosensory functions, respectively.

### Nodose ganglia clusters express both unique and overlapping sensory receptor transcripts

We next examined the cluster-defining transcripts for each of our 19 identified NG clusters (**Figure 2A**). Among these genes were a large number that encode membrane proteins previously associated with sensory functions. These include several ion channels including *Grin2a* (NMDA receptor subunit A; defining Cluster 12), *Chrna3* (nicotinic acetylcholine receptor subunit 3; defining Cluster 4), *Trpa1* (transient receptor potential cation channel subfamily A member 1; defining Cluster 3), and *Htr3b* (5-hydroxytryptomine receptor 3B; defining Cluster 5) ^28–32^. Other functional transmembrane proteins thought to be involved in visceral sensory signaling were also prominent among cluster defining genes, including *Tmem233* (defining Cluster 9)*, Cckar* (defining Cluster 4), and *Gpr65 (*G protein coupled receptor 65; Cluster 5) ^33–35^. Across all clusters, expression of the top 5 cluster-defining genes was highly correlated between right and left NG (R^2^ = 0.956, p < 0.000, Pearson’s correlation), consistent with excellent integration of data from right and left NG samples prior to clustering (**Figure 2B**).

**Figure 2.**
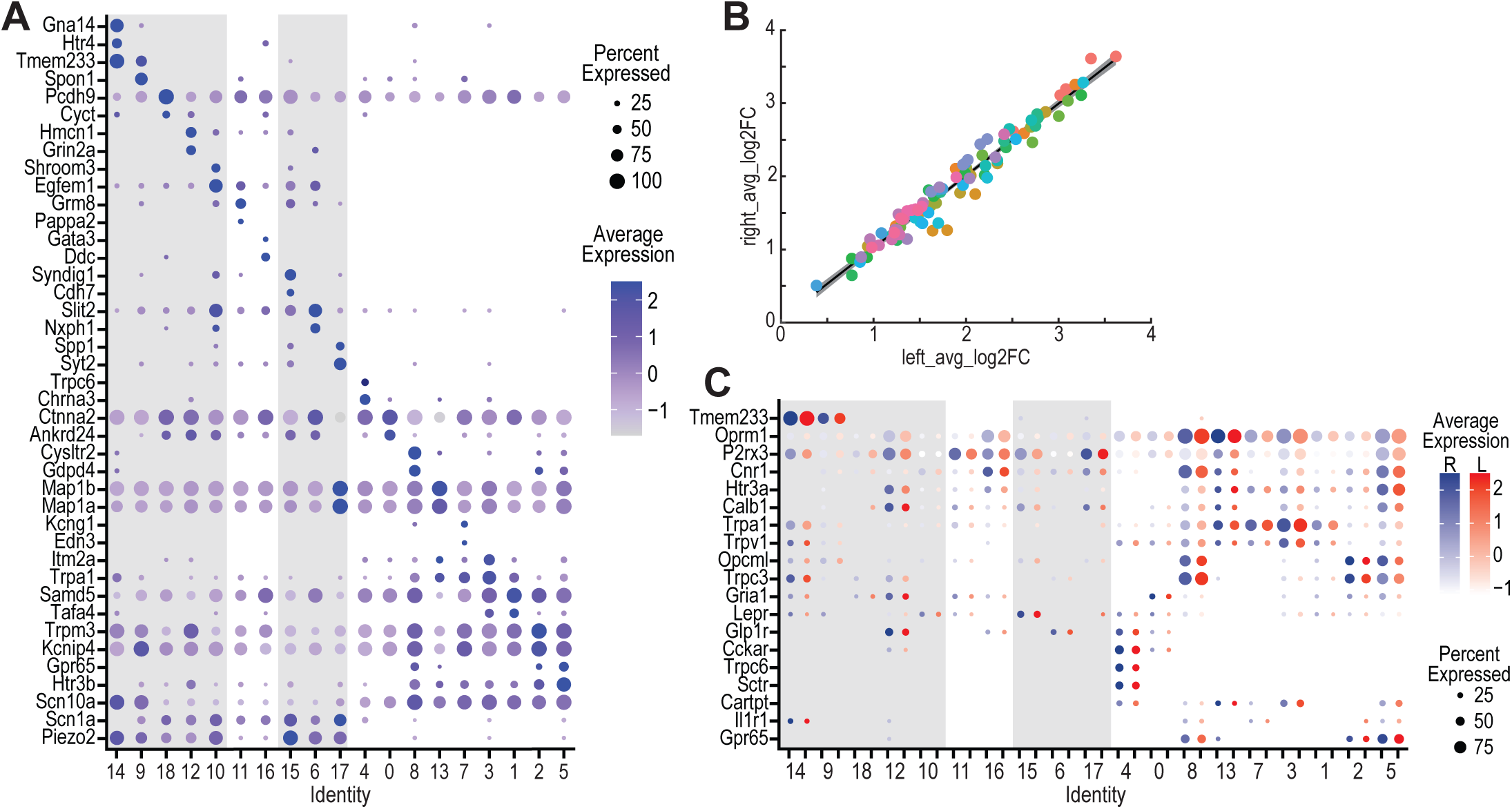
Unique genetic profiles define NG neuronal clusters. A) Expression of the top 2 cluster-defining genes for each cluster. B) Across all clusters, expression of the top 5 cluster-defining genes was highly correlated between left and right NG (R^2^ = 0.956, p < 0.0000, Pearson’s correlation), indicating strong integration of data from the two sides prior to clustering; colors represent cluster identities as in Fig. 1B. C) Expression of select genes known to be expressed in gut-innervating NG neurons. For dot plots, average expression is plotted if the gene was expressed in > 10% of cells in the cluster.

We further examined the expression of several genes previously identified in the literature to be strongly expressed in NG neurons (**Figure 2C**). Based on literature suggesting that lateralized vagal reward signaling arises from gut-innervating NG neurons ^16^, we focused on genes known to be associated with digestive signaling in the GI tract ^36–38^. However, it is worth noting that many of these receptors, including *Oprm1*, *P2rx3* (P2X purinergic receptor 3), and *Trpv1* (transient receptor potential cation channel subfamily V member 1), for example, are not exclusive to gut-innervating NG neurons, and have been shown to impact vagal signaling in multiple organ systems ^32,39–41^. Several of these known NG-expressed genes, including *Gpr65*, *Trpa1*, and *Cckar* are also included among the cluster-defining genes shown in **Figure 2A**, supporting the validity of our computational clustering approach. In this targeted subset of genes, we find that the majority are expressed to some degree in both putative mechanosensing and chemosensing clusters, though overall expression patterns support the hierarchical clustering of our subtypes along these functional lines (**Figure 2C**). Consistent with published literature, we find that expression of *Oprm1*, *P2rx3*, and *Cnr1* (endocannabinoid receptor 1), in particular, are near-ubiquitously expressed across almost all NG clusters ^42–44^

### Few genes exhibit significant left-right differential expression within NG clusters

To begin to examine the possible molecular basis of lateralized vagal reward signaling, we performed a pseudobulk analysis of differential gene expression within each of our 19 clusters ^45^. This analysis identified five clusters in which any genes were found to be significantly differentially expressed between right and left NG (Clusters 7, 3, 1, 2, and 5; **Figure 3A**) In the UMAP visualization and hierarchical clustering tree, these *Scn10a*+ chemosensing clusters were situated near each other (**Figure 1B & 1D**), suggesting a degree of similarity in their genetic profiles. Across all five clusters, 15 unique genes were identified as significantly differentially expressed with at least a 1.5-fold difference in expression between left and right sides and a Bonferroni-adjusted p < 0.01 (**Figure 3A**). Expression of two of these genes, *Gria1* (AMPA receptor subunit 1) and *Shisa9* (Shisa family member 9), was found to be significantly left-NG biased in more than one cluster. Overall, however, the effect sizes associated with identified differentially expressed genes were small, rarely exceeding a 2-fold difference in expression between sides. Our pseudobulk analysis suggests that, in naïve rats, cell-type specific expression of differential sensory receptors may not be a major driver of lateralized vagal reward signaling.

**Figure 3.**
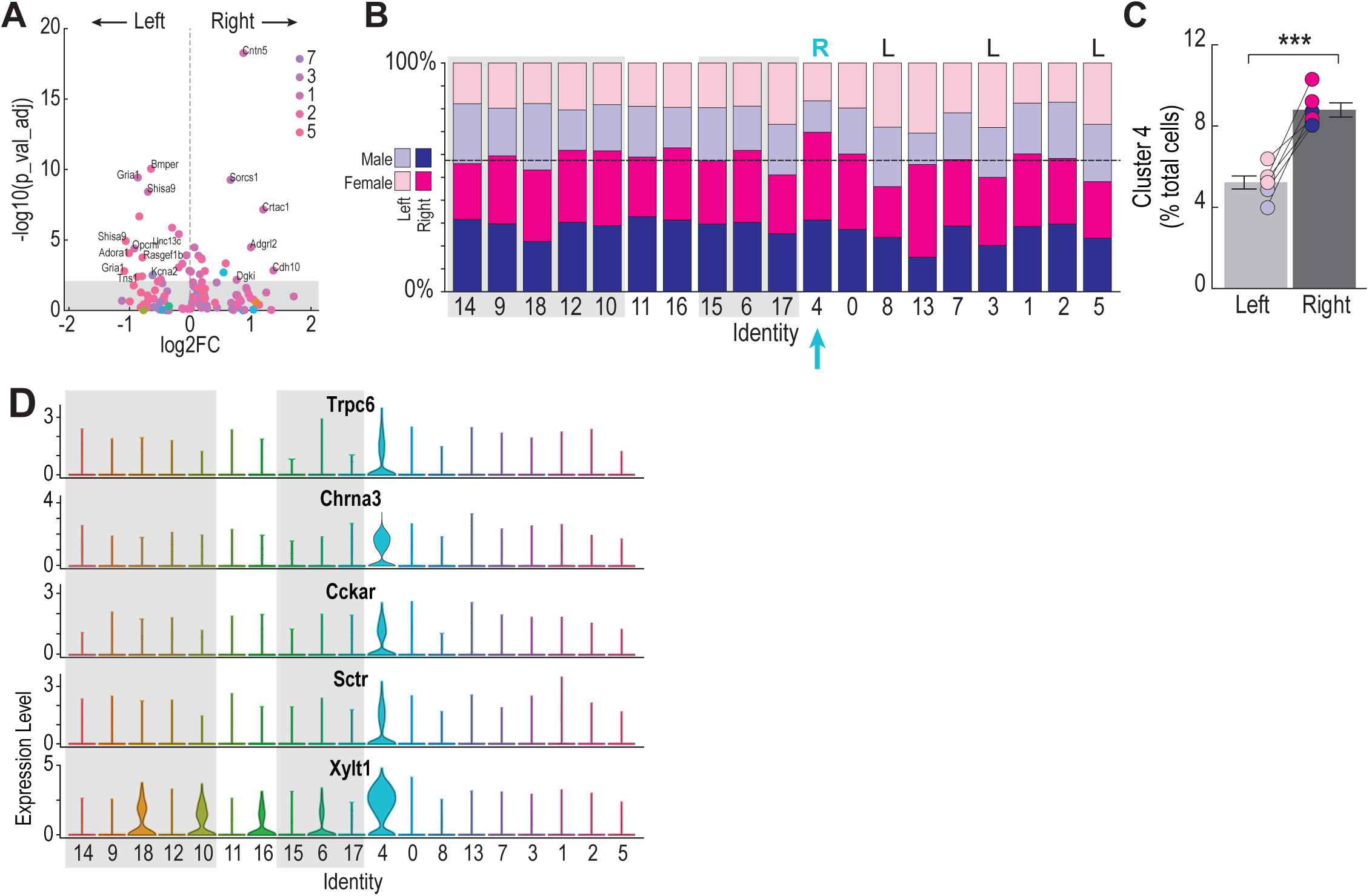
Identification of right-side biased expression of a Chrna3 and Cckar co-expressing cluster. A) Pseudobulk analysis identified 15 genes with significant left-right differential expression distributed across Clusters 7, 3, 1, 2, and 5. B) Percent of cells in each cluster originating from left (light colors) versus right (dark colors) NG samples from male (blue) vs. female (magenta) rats. Dashed line denotes percentage of total cells across all clusters obtained from right NG samples. Cluster 4 (cyan arrow) was the only cluster that exhibited significantly greater than expected representation in right NG samples. R/L above bars denotes clusters found to have significantly greater than expected representation in right (R) or left (L) NG. C) The percentage of Cluster 4 cells in left-NG vs right-NG replicate groups. Paired data represent percentage of Cluster 4 cells for each replicate from male (blue) and female (magenta) rats; error bars denote SEM; ***p < 0.001, paired t-test. D) Cluster 4 is distinguished by expression of *Trpc6*, *Chrna3*, *Cckar*, *Sctr*, and *Xylt1*.

### A *Chrna3* and *Cckar* co-expressing cluster is preferentially found in right nodose ganglia

To test whether the identified cell types themselves might be differentially expressed between right and left NG, we analyzed cell counts from each cluster to determine if that cluster contained a greater-than-expected number of cells from the left or right side. Across the entire dataset, we observed that the percentage of cells from right NG was slightly higher than the percentage of cells from left NG (56% vs. 44%, respectively), which we attribute to a slight bias in sample collection, as right NG were collected first during dissection in 20 of 36 rats. For each individual cluster, we compared the percentage of cells from the right or left NG to this expected distribution using multiple Bonferroni-corrected binomial tests. This statistical analysis revealed four clusters with significantly lateralized expression (**Figure 3B**), only one of which was a right-biased cluster. Cluster 4 comprised 7.2% of the overall NG population (n=1,694/23,428 nuclei) and made up a significantly larger percentage of right NG cells compared to left (**Figure 3C**; Left: 8.8 ± 0.30%, Right 5.2 ± 0.27%, p = 0.0006, paired t-test). This cluster was defined by high levels of expression of *Cckar* and *Sctr*—genes known to play key roles in nutrient sensing and digestive signaling ^46–48^—as well as *Trpc6* (transient receptor potential cation channel subfamily C member 6), *Chrna3, and Xylt1* (xylosyltransferase 1) (**Figure 3D**). Left-biased expression of Clusters 8, 3, and 5 was also observed. These clusters were defined by the expression of several genes involved in inflammation (e.g., *Cysltr2* and *S1pr3*, Clusters 10 and 8, respectively) and noxious stimuli (e.g., *Trpa1* and *Htr3b*, Clusters 8 & 9, respectively) known to be expressed in vagal fibers innervating the respiratory and gastrointestinal systems ^21,37,49–51^.

CCK1Rs are strongly expressed in gut-innervating NG neurons and are known to regulate satiety and macronutrient sensing ^52–55^. CCK has also been shown to play a role in the vagal signaling pathways that shape nutrient preferences ^25,56,57^. Right-biased expression of a NG population co-expressing *Cckar*, *Glp1*, and *Npy2r* was recently shown in mice (Lansbury et al., 2025), and lesions to *Cckar*+ neurons in the right, but not left NG, block the satiety-inducing effects of systemically administered CCK ^16^. Right-biased expression of a *Cckar*-expressing cluster of NG neurons thus provides a strong candidate for lateralized transduction of reward-related interoceptive signaling in rodents. To validate right-biased expression of our *Cckar*-expressing Cluster 4, we used RNA-FISH to quantify *Cckar*, *Chrna3*, and *Xylt1* expression in left versus right NG from six rats (**Figure 4A-C**). Our results confirmed overlapping expression of these 3 genes in both left and right NG cells (**Figure 4D-E**). *Cckar*+ neurons were found to be significantly more numerous in right NG compared to left (**Figure 4F**; *Cckar*+, cells/mm^2^: Left: 96.04 ± 7.64, Right: 138.94 ± 12.12; p = 0.0245, paired t-test). We also found that while most *Chrna3*+ neurons co-expressed *Cckar* and/or *Xylt1* (n = 2,513/4,309; 58.3%), 42.5% expressed *Chrna3* alone, without these other Cluster 4 marker genes (**Figure 4E**). Differential expression of *Chrna3*+ neurons exhibited a non-significant trend toward right-side biased expression (**Figure 4G**; Left: 121.62 ± 7.05, Right: 166.98 ± 12.34; p = 0.0605, paired t-test). Thus, while the majority of *Cckar*-expressing cells co-express *Chrna3* (n = 2,468/3,483; 70.9%), and this population is seen to be more numerous in right NG compared to left, we also find a population of *Chrna3*+ cells that do not express *Cckar* and do not exhibit strong differential left-right expression. *Xylt1*+ neurons were less numerous overall than predicted by the computational data, though this may in part be attributable to quantification difficulties resulting from high background staining in this probe channel. Nonetheless, the majority of *Xylt1+* neurons (n = 974/1,490; 65.4%) were found to co-express *Cckar* and *Chrna3* (**Figure 4J**), and *Xylt1*+ neurons were significantly more numerous in the right NG (**Figure 4H**; *Xylt1*+, cells/mm^2^: Left: 38.72 ± 4.08, Right: 56.67 ± 6.87; p = 0.0265, paired t-test), as were triple-labeled cells (**Figure 4I**; *Cckar*+|*Chrna3+*|*Xylt1+* co-labeled, cells/mm^2^: Left: 22.66 ± 1.02, Right: 40.99 ± 5.15; p = 0.0086, paired t-test). Overall, our RNA-FISH results confirm the presence of a *Cckar*+|*Chrna3*+|*Xylt1*+ neuronal subtype that is preferentially expressed in the right NG.

**Figure 4.**
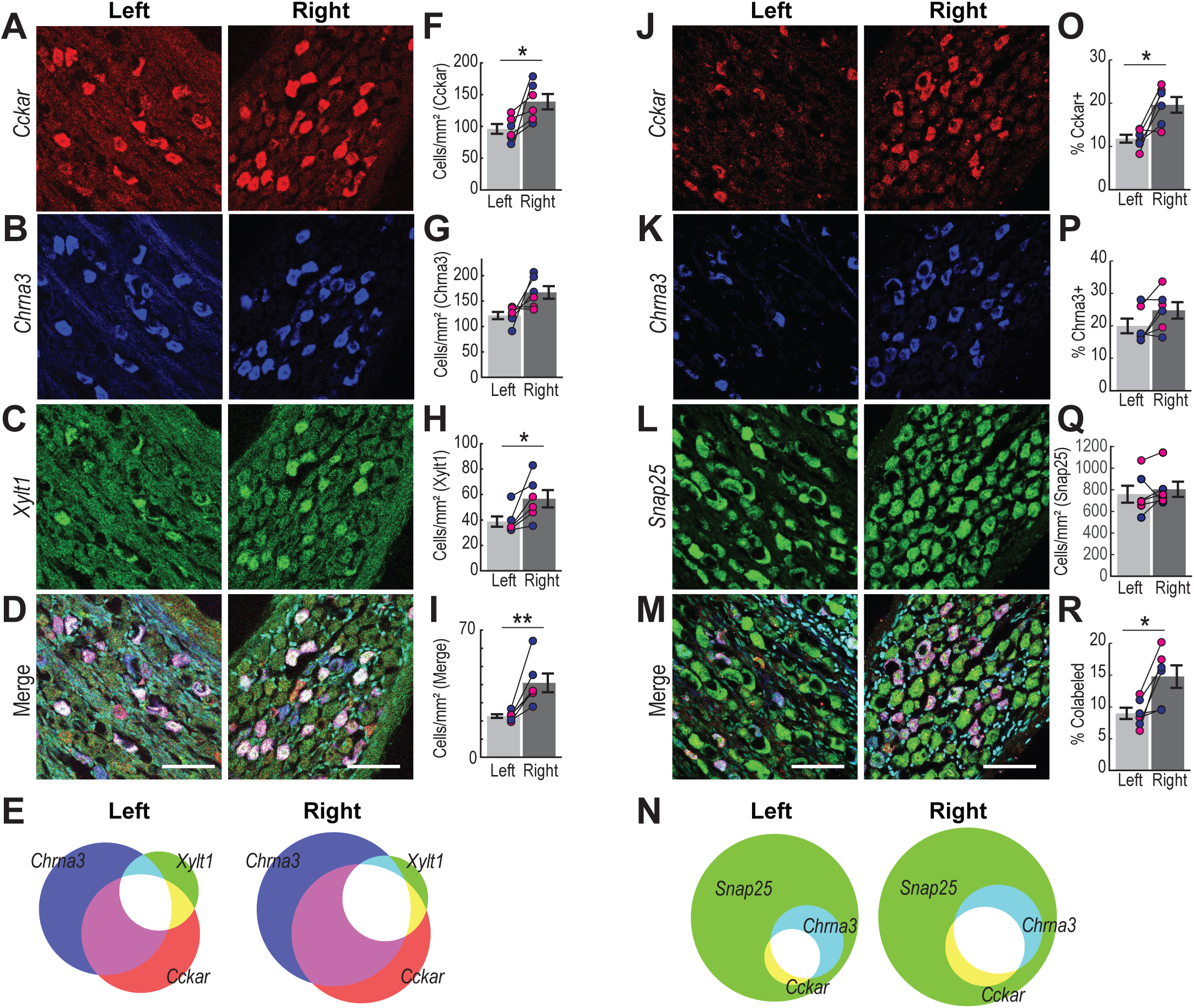
RNA-FISH validation of right-biased expression of NG neurons co-expressing Chrna3 and Cckar. A-D) RNA-FISH confirms the presence of a *Cckar* (A, red), *Chrna3* (B, blue), and *Xylt1* (C, green) co-labeled (D) population of cells in the left and right NG of rats. Merged images (D) also show DAPI-stained nuclei (cyan) and scale bar (100 µm) for A-D. E) Semi-quantitative Venn diagrams illustrating overlapping *Cckar*, *Chrna3*, and *Xylt1* labeling in left and right NG; circle area for each probe is proportional to the number of labeled cells per slice, averaged across rats. F-I) Density of cells in left vs. right NG that expressed *Cckar* (F), *Chrna3* (G), *Xylt1* (H), or were co-labeled for all 3 mRNAs (I). J-M) RNA-FISH confirms that all *Cckar*+ (J) and *Chrna3*+ (K) cells co-express the neuronal marker *Snap25* (L). Merged image (M) shows DAPI-labeled nuclei and scale bar (100 µm) for J-M. N) Semiquantitative Venn diagrams illustrating overlap of *Cckar*, *Chrna3*, and *Snap25* in left and right NG slices. O-Q) The percentage of *Cckar*+ (O) and *Chrna3*+ (P) cells in left vs. right NG, expressed as a percentage of the *Snap25*+ neuronal population (Q). R) Percentage of *Cckar*+ and *Chrna3*+ co-labeled neurons in left vs. right NG slices. Paired data in F-I and O-R represent averages across left vs. right NG slices for each rat (magenta: females; blue: males); error bars denote SEM; *p < 0.05, **p < 0.01, paired t-test.

To estimate the percentage of NG neurons comprised by our Cluster 4 subtype, we performed additional RNA-FISH experiments to probe expression of the neuronal marker *Snap25*, along with *Cckar* and *Chrna3* in left vs. right NG slices (**Figure 4J-R**). As in the previous experiment, we found significant right-biased expression of *Cckar*+ neurons (**Figure 4O**; Percent *Cckar*+: Left: 11.86 ± 0.91%, Right: 19.60 ± 1.83%; p = 0.0162, paired t-test) but differential expression of *Chrna3+* did not quite reach statistical significance (**Figure 4P**; Left: 19.97 ± 2.28%, Right: 24.77 ± 2.51%; p = 0.0661, paired t-test). The average density of cells expressing *Snap25* did not differ between right and left NG (**Figure 4Q**; *Snap25*+, cells per mm^2^: Left: 759.20 ± 78.21, Right: 804.84 ± 70.15; p = 0.4538, paired t-test). Consistent with our computational results, *Cckar* and *Chrna3* co-labeled neurons, putatively corresponding to our Cluster 4, were found to be ca. 1.7 times more numerous in the right NG compared to the left (**Figure 4R**; Percent *Cckar*+|*Chrna3*+ co-labeled: Left: 9.00 ± 0.90%, Right: 14.78 ± 1.76%, p= 0.0204, paired t-test).

### *Cckar* expression partially overlaps with expression of *Glp1r* and *Cartpt*

Right-biased co-expression of several gut peptide receptor genes, including *Cckar*, *Glp1r*, and *Npy2r*, was recently demonstrated in NG of mice (Lansbury et al., 2025), delineating lateralization of a neuronal subpopulation previously identified in transcriptomics data in that species ^21^. Our transcriptomics data from rats, however, suggests that while several genes associated with gastrointestinal peptidergic signaling, including *Glp1r*, *Cartpt*, and *Lepr*, for example, are likely present in our *Cckar*+|*Chrna3*+ Cluster 4, expression of these genes may be more prominent in *Cckar*-clusters. We thus performed additional RNA-FISH experiments to quantify whether a subset of these genes, namely *Glp1r* and *Cartpt*, also exhibit differential expression in left vs. right NG in rats (**Figure 5A-D**), and the extent to which these genes are co-expressed in *Cckar*+ NG cells (**Figure 5E**). Our results again confirm strong right-biased *Cckar* expression (**Figure 5F**; *Cckar*+, cells/mm^2^: Left: 75.35 ± 13.11, Right: 131.68 ± 16.40; p = 0.0202, paired t-test). While the majority of *Cckar*+ cells were found to co-express *Cartpt* (n = 2,213/3,250, 68.1%), the reverse was not true—most *Cartpt*+ cells did not co-express *Cckar* or *Glp1r* (**Figure 5E**; n=6,866/9,387, 73.1%), and differential expression of *Cartpt*+ cells did not reach statistical significance (**Figure 5G**; Left: 255.04 ± 24.39, Right: 316.09 ± 15.23; p = 0.0598, paired t-test). Consistent with our computational results, and similar to what has recently been reported for mice ^12^, just over half of *Cckar*+ cells were found to co-express *Glp1r* (n = 1,720/3,250, 52.9%). *Glp1r*+ cells were found to be significantly more numerous in right NG compared to left (**Figure 5H**; Left: 92.69 ± 5.64, Right: 134.12 ± 10.38; p = 0.0321, paired t-test), as were triple-labeled cells (**Figure 5I**; Left: 31.05 ± 5.64, Right: 51.44 ± 8.93; p = 0.0296, paired t-test). These results indicate that *Cartpt* expression labels a much broader population of NG neurons than *Cckar* and *Chrna3* co-expression. By contrast, there is significant overlap and similar right-biased expression in *Glp1r+* and *Cckar*+|*Chrna3*+ NG neuron populations in rats.

**Figure 5.**
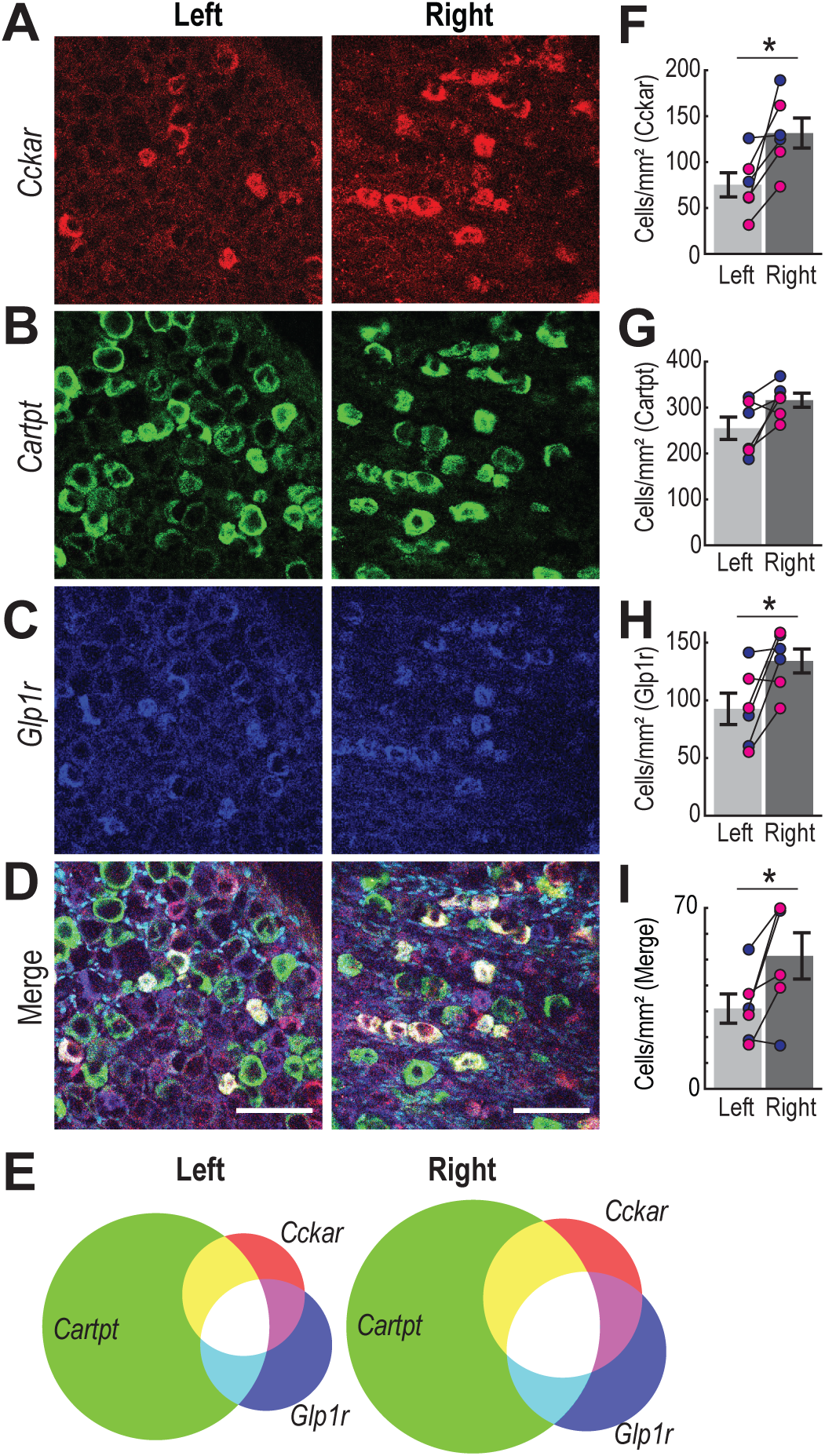
Cckar+ NG neurons overlap with populations expressing Cartpt and Glp1r. A-D) RNA-FISH was used to probe for overlapping expression and right-side biased expression of *Cckar*+ (A, red), *Cartpt*+ (B, green), *Glp1r*+ (C, blue), and triple-labeled neurons (D) in left and right NG slices from rats. Merged images (D) also show DAPI-stained nuclei (cyan) and scale bar (100 µm) for A-D. E) Semiquantitative Venn diagrams, as in Figure 4E, illustrating overlap in *Cckar*, *Glp1r*, and *Cartpt* labeling in left and right NG. F-I) Density of cells in left vs. right NG that expressed *Cckar* (F), *Cartpt* (G), *Glp1r* (H), or were co-labeled for all 3 mRNAs (I). Paired data represent averages across left vs. right NG slices for each rat (magenta: females; blue: males); error bars denote SEM; *p < 0.05, paired t-test.

### *Cckar* and *Chrna3* co-expressing NG neurons exhibit right-biased expression in mice

By integrating our rat data with a publicly available dataset from mice ^58^, we examined whether gene expression patterns and right-side bias of a *Cckar*+ cluster could be similarly observed in both species. Mouse and rat datasets were first filtered using quality control metrics (**Figure S3A-B)**. After QC filtering, the mouse and rat datasets were integrated and clustered, applying the same computational pipeline as before. Clusters representing jugular neurons (*Pdrm12*+) and non-neuronal cell types (expressing *Ttn*, *Myh4*, *Mpz*, *Mbp*, *Hand2*, *Daam2*, and/or *Mecom*) were removed through 3 rounds of initial clustering (**Figure S4**). After removal of non-neuronal and jugular neuron clusters and subsequent reclustering, 16 putative NG clusters were identified in the integrated dataset (**Figure 6A**), with good overlap between the mouse and rat data (**Figures 6B-C**). In the integrated dataset, Cluster 7 was observed to be defined by expression of *Trpc6* and *Chrna3*, and to express high levels of *Cckar*, similar to Cluster 4 in our rat-only data (**Figure 6D**). In both mice and rats, Cluster 7 made up ca. 5-6% of the total sample (mouse: n=181/3,664 nuclei, 4.9%; rat: n=1,766/29,117 nuclei, 6.1%). Statistical analysis confirmed that neurons in this cluster were significantly more numerous in right NG samples than in left in both mice (n = 169/181, 93%, compared to 62.5% right NG neurons total in mouse samples; p < 0.0000, binomial test), and rats (n = 1,228/1,766; 69.5%, compared to 59.7% right NG neurons total in rat samples; p < 0.0000, binomial test). While *Chrna3* and *Cckar* co-expression defined integrated Cluster 7 and exhibited strong expression in both the mouse and rat datasets (**Figure 6E; Figure S3C**), *Xylt1* appeared weakly expressed within this cluster in mouse samples. Several other well-known NG genes associated with gut-brain signaling, including *Cartpt*, *Npy2r*, and *Vip*, were observed to be much more strongly expressed in Cluster 7 in mice compared to rats (**Figure 6E**), suggesting significant species specificity in gene expression patterns within this cluster.

**Figure 6.**
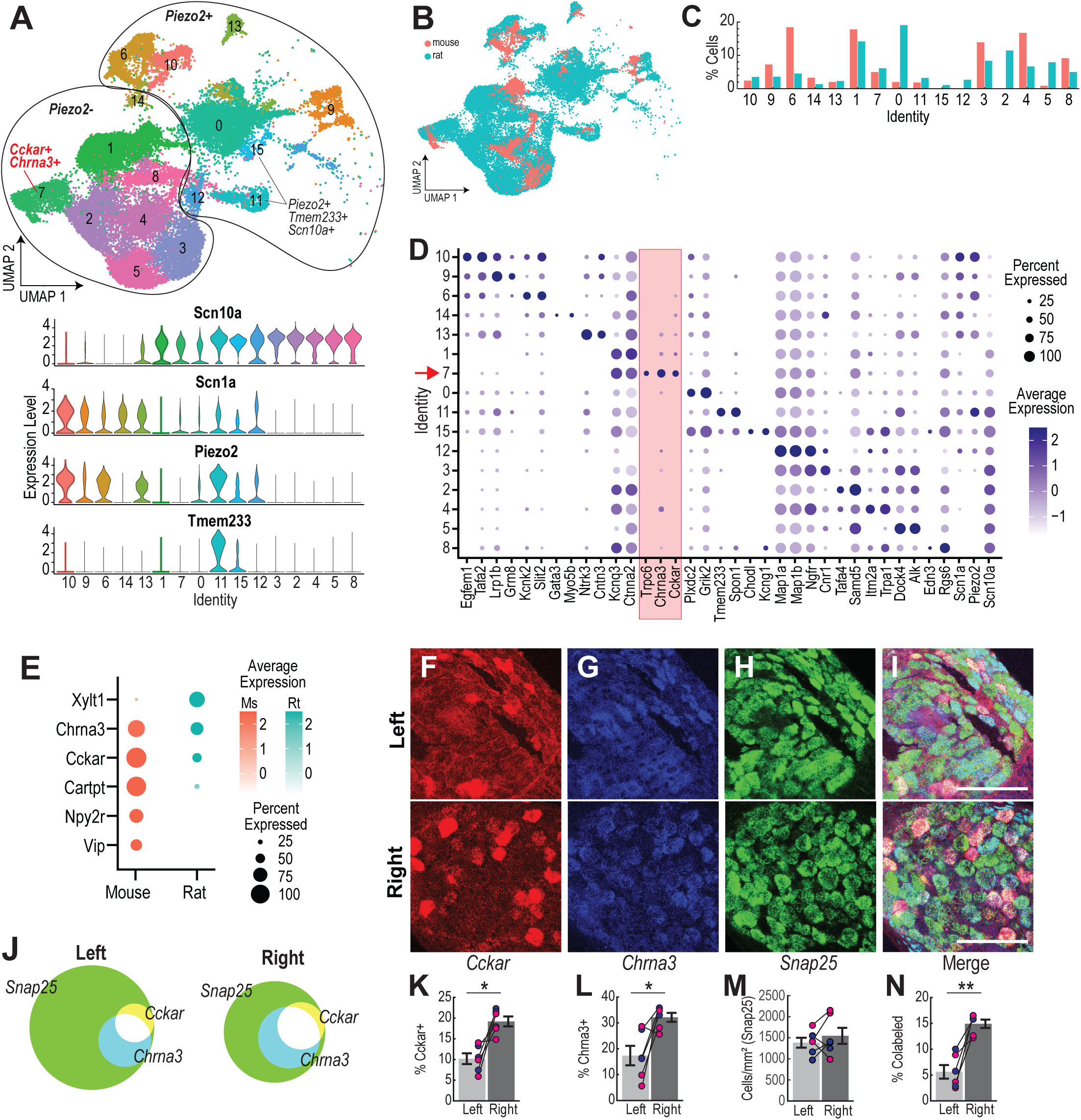
Mice also exhibit preferential expression of a Chrna3+ and Cckar+ subtype in right NG. A) Clustering identified 16 putative neuronal clusters in the integrated rat + mouse snRNA-seq dataset (top). Putative mechanosensitive versus chemosensitive clusters can be distinguished based on expression of *Piezo2*. As in rats, *Piezo2* expression overlaps with expression of *Scn1a* in most mechanosensory clusters, whereas most *Scn10a+* clusters in the integrated dataset are *Piezo2*-(bottom). B-C) Overlapping UMAP representations (B) and percentages of cells from each dataset distributed across identified clusters (C) indicate rat and mouse data were well integrated. D) Dot plot illustrating expression of top 2 marker genes for each cluster, plus *Cckar* expression across all clusters. Cluster 7, defined by high expression of *Trpc6* and *Chrna3*, and expressing high levels of *Cckar*, is highlighted (red arrow and shading). E) In both mouse and rat, co-expression of *Chrna3* and *Cckar* is predicted within Cluster 7, though expression of other marker genes (*Xylt1*) and digestive peptide signaling related genes (*Cartpt*, *Npy2r*, and *Vip*) diverges between the two species. F-I) RNA-FISH was used to quantify *Cckar* (F), *Chrna3* (G), and *Snap25* (H) expression, in left (top) and right (bottom) NG slices from mice; DAPI staining and scale bar (100 um) for F-I is shown in merged images (I). J) Semiquantitative Venn diagrams, as in Figure 4E, illustrating overlap in *Snap25*, *Cckar*, and *Chrna3* labeling in mouse NG slices. K-M) Percentage of *Cckar*+ (K) and *Chrna3*+ (L) cells in left vs. right NG, expressed as a percentage of the *Snap25*+ neuronal population (M). N) Percentage of *Cckar*+ and *Chrna3*+ co-labeled neurons in left vs. right NG slices. Paired data in K-N represent averages across left vs. right NG slices for each mouse (magenta: females; blue: males); error bars denote SEM; *p < 0.05, **p < 0.01, paired t-test.

To validate right-biased expression of our integrated Cluster 7 in mice, we performed RNA-FISH to probe for *Cckar*, *Chrna3,* and *Snap25* expression in mouse NG slices (**Figure 6F-N**). As in rats, most *Cckar*+ neurons in mice also co-expressed *Chrna3* (**Figure 6J**; n = 763/1,058, 72.1%), and *Cckar*+ neurons were found to be significantly more numerous in right NG compared to left (**Figure 6K**; Left: 10.18 ± 1.30%, Right: 19.20 ± 1.19%; p = 0.0120, paired t-test). Only ca. 40% of *Chrna3* neurons were found to co-express *Cckar* (n = 763/1,821, 41.9%). The overall *Chrna3*+ population was found to be significantly larger in right NG (**Figure 6L**; Left: 17.35 ± 3.77%, Right: 32.14 ± 1.74%; p = 0.0214, paired t-test) as was the *Cckar*+|*Chrna3*+ co-labeled population (**Figure 6N**; Left: 5.62 ± 1.30%, Right: 14.91 ± 0.81%; p = 0.0024, paired t-test). We found no difference in *Snap25*+ neuron density between right and left NG (**Figure 6M**; *Snap25*+, cells per mm^2^: Left: 1,386.22 ± 117.63, Right: 1,548.02 ± 191.49; p = 0.4030, paired t-test).

Combined, our results demonstrate that a *Cckar* and *Chrna3* co-expressing subpopulation of neurons is preferentially expressed in the right NG over the left in both mice and rats.

### CCK1R agonism preferentially activates right NG neurons

To validate functional lateralization of CCK1R signaling in the NG, we performed calcium imaging on left and right NG cultures from 10 rats (6 male, 4 female). Responses to bath application of the selective CCK1R agonist, A71623 (100 nM), were recorded for 1,233 total neurons (**Figure 7A-B**). Consistent with our transcriptomic and RNA-FISH results, we observed significant asymmetry in population responsiveness to CCK1R agonism among left vs. right NG neurons. Approximately twice as many A71623-responsive neurons were found in right NG cultures compared to left (**Figure 7C**; Right: n=69/495, 13.94%; Left: n = 50/738, 6.78%; p < 0.0001, Chi-square test). Among responders, there was no difference between left and right NG neurons in the magnitude of the peak response to CCK1R agonism (**Figure 7D**; Left: 49.41 ± 3.68%, Right: 45.42 ± 2.03%, p = 0.3117, unpaired t-test). These findings are consistent with our computational data and suggest that protein expression levels are similar for CCK1R-expressing left and right NG neurons. However, the CCK1R-expressing population itself is more numerous in the right NG compared to the left.

**Figure 7.**
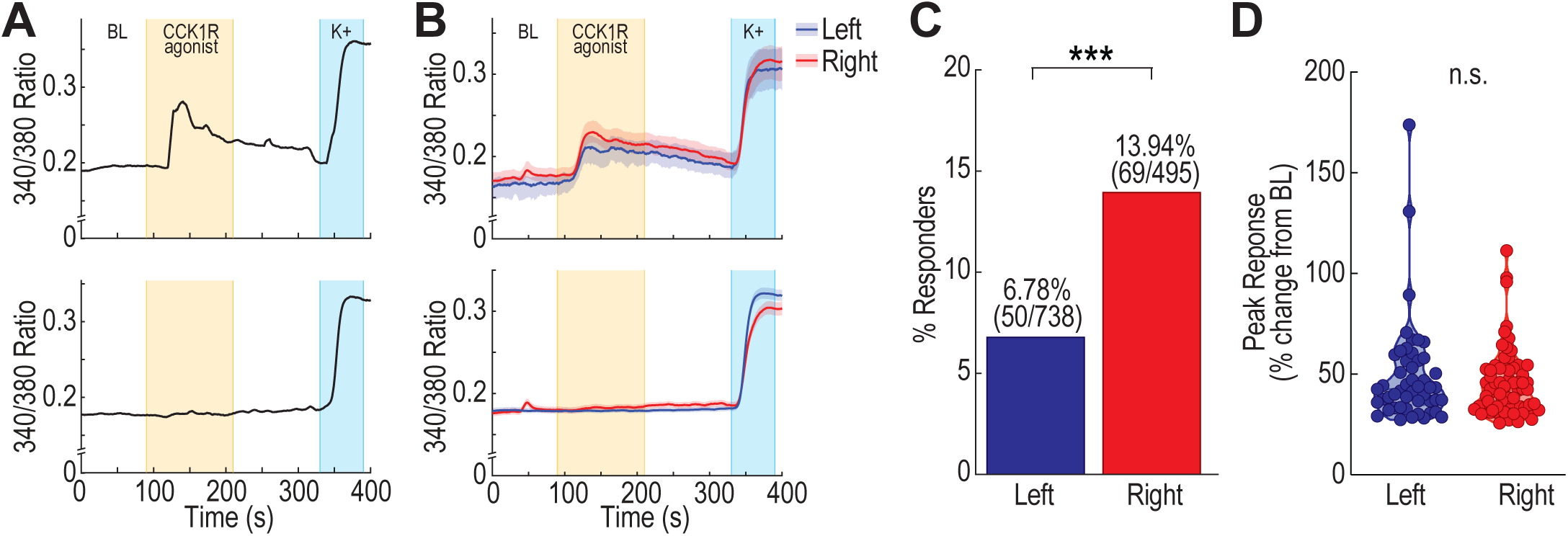
Right-biased responses to CCK1R activation in rat NG. In vitro calcium imaging was performed on primary cultures from left and right NG of rats during bath application of the CCK1R agonist, A71623. A) Example traces from neurons classified as A71623-responsive (top) or non-responsive (bottom) based on the presence or absence of an increase in calcium signal following drug application. B) Mean calcium signals of A71623-responsive (top) and non-nonresponsive (bottom) neurons from the left (blue) and right (red) NG. Shading represents 95% confidence intervals. C) Significantly more A71623-responsive neurons were found in right-NG cultures than in left-NG cultures; ***p < 0.001, Chi-square. D) Among neurons exhibiting A71623 responses, the peak amplitude of the calcium signal did not differ between left and right NG samples.

Taken together, our transcriptomic, RNA-FISH, and calcium imaging results demonstrate significant right > left asymmetry in the expression of a *Cckar*+ vagal neuron subtype, and in functional responsiveness to CCK1R activation in the NG.

## DISCUSSION

Our data represent the first snRNA-seq analysis of NG neurons in rats, providing an important resource to the neuroscientific community. In the current study, we aimed to utilize this dataset to determine whether genetic differences in right vs. left NG neurons could contribute to right-biased encoding of visceral reward-related signaling by the vagus nerve. Our analyses identified a single neuronal cluster, defined by *Cckar* and *Chrna3* co-expression, that exhibits preferential expression in the right NG of both rats and mice. Our transcriptomic and RNA-FISH results further indicate that, in rats, neurons in this population are further defined by *Sctr* co-expression, and often co-express *Cartpt* and *Glp1r*, pointing to a role for this neuronal subtype in digestive signaling. Finally, our calcium imaging experiments provide novel evidence of functional right-biased asymmetry in CCK1R-mediated NG signaling. Combined, the current study identifies a unique population of right-biased *Cckar* and *Chrna3* co-expressing NG neurons as a strong candidate for contributing to the lateralized vagal encoding of interoceptive rewards.

Cholecystokinin (CCK) is a peptide released by enteroendocrine cells in the gastrointestinal tract in response to food intake, and CCK1Rs are strongly expressed in gut-innervating NG neurons ^52–54^. CCK1Rs play an important role in regulating satiety and sensing macronutrients through gut-brain vagal signaling pathways. CCK has also been found to play a critical role in shaping the preference for macronutrients including fat, sugar, and amino acids ^25,56,57^. Glucagon-like peptide-1 (GLP1) and secretin are likewise known to contribute to vagal-dependent nutrient detection, satiety signaling, and the modulation of reward-related signaling pathways ^59–62^. In light of this prior work, our data support a model of sensory transduction of reward-related interoceptive cues by a right-side biased population of neurons engaged in detection of CCK and other digestive peptides.

Previous studies have shown that stimulation of the right, but not left, vagus nerve increases dopamine release and reinforces behavior, consistent with lateralization of interoceptive reward processing ^16,18^. Gut-innervating vagal neurons, in particular, likely give rise to this left-right asymmetry in vagal-mediated reward signaling ^16^. Using an unbiased transcriptomic approach, we identified a single cluster of NG neurons, distinguished by co-expression of *Cckar* and *Chrna3* in both rats and mice, that is significantly more abundant in the right NG compared to the left. Our transcriptomic data indicate that this neuronal subtype is also distinguished by high levels of expression *Trpc6* and *Sctr*. This gene expression profile is highly similar to that of a putative gut-innervating IGLE population previously identified by Kupari et al. ^21^ in mice. Our study provides novel evidence that this cluster is conserved across mice and rats and that it exhibits strongly right-biased expression that is unique among NG subtypes.

Expression of this *Cckar* and *Chrna3* co-expressing subpopulation is ca. 1.5 to 2 times greater in the right NG compared to the left, comprising up to ca. 20% of right NG neurons in our data, and ca. 5-10% of left NG neurons. It is notable, then, that VNS has been observed to exhibit rewarding properties only when applied to the right side. Several factors may contribute to these findings. At parameters typically used for neurological indications, VNS is thought to preferentially activate myelinated cervical vagal fibers ^63^, and thus may only weakly activate *Cckar*+ IGLEs ^37^. Clinically and preclinically, VNS is predominantly applied to the left vagus nerve, which our results indicate is likely to harbor fewer of these *Cckar*+ gut-innervating, putative reward-transducing neurons. As a result, left-sided stimulation may fail to sufficiently engage this population, and/or the effects of activation may be masked by co-activation of mechanosensitive or nociceptive-like subtypes. On the right side, the impact of stimulation of the larger population of vagal reward-sensitive neurons may be further amplified by asymmetrical connectivity within the central nervous system, as has been described by Han et al. ^16^, producing dopaminergic signaling that more readily drives behavioral reinforcement. The current study presents a novel characterization of prominent neural asymmetry in left vs. right vagal signaling and highlights the importance of considering this molecular and functional lateralization in future mechanistic studies of interoceptive reward processing. By demonstrating significant differential left-right expression of neuronal subtypes in the NG, our work additionally provides fundamental insights to inform the development and optimization of VNS or pharmacological strategies designed to modulate vagal signaling. These insights may be particularly relevant for therapeutic strategies aimed at treating conditions involving dopamine dysregulation, including obesity, depression, and addiction.

## Limitations of the study

In this study, we investigated whether genetic differences in right vs. left NG neurons could contribute to right-biased encoding of visceral reward-related signaling by the vagus nerve. We identified a single cluster, marked by the expression of *Cckar* and *Chrna3*, that exhibits a strong right-biased expression in the NG. Combined with prior literature, our results suggest that lateralized reward-related interoceptive signaling may be supported by this unique neuronal subtype. Our snRNA-seq data indicate that this *Cckar*+ and *Chrna3*+ population co-expresses genes encoding several digestive peptide receptors, consistent with a gut-innervating subtype. However, the extent to which the gut receives preferential innervation from *Cckar*+ neurons in the right NG rather than the left, remains to be anatomically confirmed. Additional research is also needed to causally test whether activation of this right-biased NG subpopulation is indeed sufficient to drive dopaminergic midbrain activity and reinforce behavior, as well as to determine the specific cellular signaling pathways that participate in the lateralized transduction of visceral rewards by the vagus nerve.

## ACKNOWLEDGMENTS

This work was funded by NIH R21 DA055166 (C.A.T.), R01 NS065926 (T.J.P.), R01 DK115478 (B.J.K.), and F31 NS129269 (V.M.H.). We thank Dr. Adriana Shembel, Dr. Robert Morrison, and Dr. I-Fan Mau for their technical assistance in establishing NG dissection protocols. We thank the UTD Imaging and Histology Core for their expertise in imaging methods. We thank Dr. Ashley Plumb and Dr. Katelyn Sadler for their assistance with calcium imaging experiments.

## AUTHOR CONTRIBUTIONS

H.F.W. participated in conceptualization, investigation, formal analysis, visualization, data curation, and writing – original draft; V.M.H., I.S., H.Q., and K.M. participated in investigation and writing – review & editing; B.J.K. and T.J.P. participated in supervision, writing – review & editing, and funding acquisition; C.A.T. participated in conceptualization, formal analysis, data curation, writing – original draft, project administration, and funding acquisition.

## DECLARATION OF INTERESTS

The authors declare no competing interests.

## METHODS

### EXPERIMENTAL MODEL DETAILS

A total of 52 rats (27 male, 25 female) and 6 mice (3 male, 3 female) were used in the study. All procedures were conducted in accordance with the National Institutes of Health Guide for the Care and Use of Laboratory Animals and were approved by the University of Texas Institutional Animal Care and Use Committee (protocol number 2024-0126).

### METHOD DETAILS

#### Experimental design

The right and left NG of experimentally naïve, 8-week-old male (n=18) and female (n=18) rats were dissected, and single-nucleus RNA sequencing (snRNA-Seq) was performed. snRNA-seq was performed on NG samples from 12 replicate groups (3 Male|Left, 3 Male|Right, 3 Female|Left, 3 Female|Right); each replicate group was created by pooling left or right NG samples from six rats (**Figure 1A**). Samples were prepared using a Parse Biosciences Evercode Nuclei Fixation Kit v2 (ECF2003) and Parse Single Cell Whole Transcriptome Kit v2 (ECW02130) and sequenced on an Illumina Nextseq 2000 at the Genome Center at the University of Texas at Dallas. Clustering and statistical analyses were performed in R version 4.4.1 ^65^ using Seurat version 4.3.0 (Hao et al., 2021). Computational results were validated using RNA fluorescence *in situ* hybridization (RNA-FISH) to quantify overlap in expression of key transcripts as well as side-specific expression in NG samples collected from an additional 6 rats (3 male, 3 female) and 6 mice (3 male, 3 female). Calcium imaging was performed on cultured NG cells from an additional 10 rats (6 male, 4 female) to validate functional asymmetry in NG CCK1R signaling.

#### snRNA-Seq Sample Preparation and Sequencing

Rats were deeply anesthetized with 5% isoflurane and sacrificed via rapid decapitation. The right and left NG were dissected and stored in separate PCR tubes before flash-freezing on dry ice. NG samples were stored in a -80 ⁰C freezer until sample preparation.

Individual NG were pooled to prepare three replicate groups for each of the four treatment groups (Male|Left; Male|Right; Female|Left; Female|Right). Each replicate group was created by pooling the right or left NG samples from 6 rats (**Figure 1A**). Nuclei were fixed and barcoded according to Parse Biosciences protocols (Evercode Version 2 Nuclei Fixation Kit, Version 2 WT Kit). The samples were split into 8 sublibraries and were sequenced via paired-end sequencing using the Illumina Nextseq 2000 at the Genome Center at the University of Texas at Dallas. In all, an estimated 90,000-100,000 nuclei were prepared for sequencing.

#### snRNA-Seq Analysis

The 12 replicate groups generated a total of 36,741 sequenced cells with an average of ca. 32,000 reads/cell and ca. 1,500 genes detected per cell. Cells were filtered using standard quality control metrics (less than 6,500 genes per cell, less than 20,000 molecules per cell, and less than 5% mitochondrial genes), removing 281 poor-quality cells. Data from each treatment group was integrated, scaled, and clustered using default parameters in Seurat. Additional filtering was performed to remove clusters expressing high levels of marker genes for muscle (*Ttn*, *Myh4*), myelin (*Mbp*, *Mpz*), endothelial cells (*Emcn*), melanocytes (*Tryp1*), satellite glia (*Apoe*, *Sparc*), and jugular neurons (*Prdm12*). This iterative process of removing clusters, reclustering, and re-examining non-neuronal cell and jugular neuron marker gene expression in the rat dataset was performed over 4 initial rounds of cluster filtering (**Figure S2**). The resulting dataset of putative NG-specific neurons was comprised of 23,428 nuclei and was reclustered once more prior to analysis.

Cluster-specific marker genes were identified using Wilcoxon rank-sum tests with Bonferroni corrections. To test for differential gene expression between right and left samples, pseudobulk analysis was used ^45^. For each cluster, data from all replicate groups was aggregated and differential right vs. left gene expression was calculated using Bonferroni-corrected Wilcoxon rank-sum tests.

To compare the lateralized expression of NG clusters in rats and mice, we integrated our rat snRNA-seq data with a publicly available mouse NG sequencing dataset of 5,507 cells in which left and right samples were separately identified (GEO #GSE185173) ^25^. The rat and mouse datasets were individually filtered using standard quality control metric thresholds (less than 6,500 genes per cell, less than 20,000 molecules per cell, and less than 5% mitochondrial genes). Mouse and rat data were then integrated, scaled, and clustered using default parameters in Seurat. Clusters in the integrated dataset were further filtered to restrict analysis to putative NG neurons by removing clusters expressing high levels of marker genes for muscle (*Ttn*, *Myh4*), myelin (*Mbp*, *Mpz*), sympathetic neurons (*Hand2*), oligodendrocytes (*Daam2*), endothelial cells (*Mecom*), and jugular neurons (*Prdm12*). There were high levels of *Apoe* across multiple clusters in the mouse data, which could be a general sign of cellular damage or stress ^69,70^, rather than a reliable marker of satellite glial cells. Thus, we did not remove *Apoe*+ clusters in the integrated dataset as we did for the rat-only data. This iterative process of removing clusters, reclustering, and re-examining non-neuronal cell and jugular neuron marker gene expression in the integrated dataset was performed over 3 initial rounds of cluster filtering (**Figure S4**). Cluster filtering resulted in an integrated dataset comprised of 32,781 nuclei (mouse: n = 3,664; rat: n = 29,117).

#### In situ hybridization

Rats and mice were deeply anesthetized with sodium pentobarbital and transcardially perfused with cold phosphate buffered saline (PBS) followed by 4% paraformaldehyde (PFA) in PBS. NG were dissected, embedded in OCT, quickly frozen over dry ice and stored at -80 ⁰C until sectioning. Slices were made at 20 µm thickness using a Leica CN1860 Cryostat and directly mounted on slides. Slides were stored at -80 ⁰C until RNA-FISH was performed.

Hybridization chain reaction RNA fluorescence *in situ* hybridization (HCR RNA-FISH v3.0; Molecular Instruments) was used to validate differential right vs. left NG expression of the computationally identified right-biased *Chrna3*+ and *Cckar*+ cluster. Hybridization probes sets targeting rat *Chrna3*, *Cckar*, and *Xylt1* were custom designed based on the transcript sequences found in NCBI (NCBI Accession numbers: *Cckar* (rat): NM_012688, *Chrna3* (rat): NM_052805, *Xylt1* (rat): NM_022295). To validate neuronal expression of the *Cckar* and *Chrna3* co-expressing populations in rats and mice, probe sets targeting *Cckar* and *Chrna3*, as well as the neuronal marker *Snap25*, were used (NCBI Accession numbers: *Snap25* (rat): NM_030991; *Cckar* (mouse): NM_009827, *Chrna3* (mouse): NM_145129, *Snap25* (mouse): NM_011428). To quantify overlapping and right-biased expression of select genes associated with digestive peptide signaling in rats, probes targeting *Cartpt*, *Glp1r*, and *Cckar* were used (NCBI Accession numbers: *Cartpt* (rat): NM_017110, *Glp1r* (rat): NM_012728). Multiplexed staining was performed using an HCR RNA-FISH kit including three-probe sets, probe channel-specific HCR amplifiers (each consisting of a pair of fluorophore-tethered hairpins), as well as hybridization, wash, and amplification buffers.

HCR^TM^ RNA-FISH was performed according to adapted vendor protocols ^64^. After removing samples from -80°C, mouse slides were baked for 30 minutes at 60°C using HybEZ II oven (#240200, ACD) to improve tissue adhesion (rat slides were not baked). The procedure for mouse samples continued as written. Slides were immersed at room temperature in 100% ethanol for 15 minutes, followed by immersion in 100% ethanol for 5 minutes, then immersed once in 1X PBS briefly to wash off remaining OCT. Slides were then air dried and a hydrophobic barrier was drawn around the tissue using the ImmEdge^®^ Hydrophobic Barrier PAP pen (H-4000, Vector Laboratories). Probe hybridization buffer was carefully added on top of the samples and allowed to pre-hybridize for 10 minutes in a humidified chamber using HybEZ II oven at 37 °C. A probe solution was prepared by adding 2.5 µL of each hybridization probe set (1 µM stock) in 100 µL of pre-heated probe hybridization buffer solution at 37 °C per slide. Pre-hybridization solution was removed from slides, and probe solution was added on top of the samples and incubated overnight (> 20 hrs) at 37 °C. As a negative control, one slide was treated with hybridization solution with no gene probes so there would be no initiator to trigger growth of the fluorescent amplification polymer.

The following day, slides were washed in a series of mixed and pre-heated probe wash buffer and 5X Saline Sodium Citrate buffer (SSCT) (BP1325, Fisher) solutions at 37 °C for 15 min each in the following order: (1) 75% probe wash buffer with 25% 5X SSCT, (2) 50% probe wash buffer with 50% 5X SSCT, (3) 25% probe wash buffer with 75% 5X SSCT, and (4) 100% 5X SSCT. This was followed by a 5 min wash in 100% 5X SSCT at room temperature. Amplification buffer was added on top of the samples, and slides were pre-amplified in the humidified chamber for 30 minutes at room temperature. Hairpin dyes were snap-cooled by incubating at 95°C for 90 seconds, then immediately cooled at room temperature in the dark for at least 30 minutes. The snap-cooled hairpins were added to the amplification buffer to create a hairpin solution. The pre-amplification buffer was removed from the slides, and the hairpin solution was added on top of the samples and incubated overnight in a humidified chamber at room temperature.

The next day, slides were washed in 100% 5X SSCT twice for 30 min and then once for 5 min. An autofluorescence quenching solution was created by making a 1:1:1 solution of reagents in the Vector^®^ TrueVIEW^®^ Autofluorescence Quenching Kit (SP-8400-15, Vector Laboratories). This solution was added on top of the samples and incubated for 3.5 minutes, then slides were washed in 1X PBS for 5 minutes at room temperature. Slides were cover slipped with VectaShield^®^ Vibrance^®^ Antifade Mounting Medium with DAPI (H-1800-10, Vector Laboratories) and sealed with nail polish. Slides were imaged within 24 hours.

#### FISH imaging and image analysis

An Olympus FV3000RS or Evident FV4000 confocal microscope was used to capture images. The same acquisition parameters were used for all experimental samples and negative controls. Two to six slices per ganglia were imaged at 20X (rat) or 60X (mouse) for semi-automated cell counting. Multiple 16-bit images with overlapping fields of view were taken to capture the entirety of each slice, and images were stitched together to reconstruct whole-slice images in each channel ^71^. For some images, the built-in multi-area time lapse (MATL) feature was used to stitch whole images. Image analysis was performed using FIJI ^67^. Image quantification was performed by a blinded experimenter. Stitched images were smoothed using a 3x3 mean filter, then thresholded using minimum pixel intensity cutoffs based on probe-specific signal-to-noise ratios (*Cckar* (rat): Mean+2.5*STD, *Chrna3* (rat): Mean+1.5*STD, *Xylt1* (rat): Mean+2.5*STD, *Snap25* (rat): Mean+1.5*STD, *Cartpt* (rat): Mean+1*STD, *Glp1r* (rat): Mean+2*STD; *Cckar* (mouse): Mean+1*STD, *Chrna3* (mouse): Mean+1.5*STD, *Snap25* (mouse): Mean+1*STD). Cells were then automatically detected using particle analysis for particles larger than 50 pixels^2^. This process generated image masks comprised of detected cells, from which cell counts for each individual channel were obtained. For each slice, probe co-expression was manually quantified by an experimenter blinded to sample side and sex, using overlaid and dual-or tri-color channel masks. Single and co-labeled cell counts for each slice were normalized by dividing by the area of the slice (mm^2^), and these normalized values were averaged across slices to obtain an average cell density for each animal. For slices with *Snap25* label, the numbers of *Cckar*+, *Chrna3*+, and *Cckar*+|*Chrna3*+ co-labeled cells were divided by the number of *Snap25*+ neurons for each slice, and these percentages were averaged across slices for each animal. To illustrate the overlap in probe labeling for each probe set, semi-quantitative Venn diagrams were created using DeepVenn ^68^. Circle area in the diagrams is proportional to the average number of labeled cells per slice in the relevant probe channel. This average was calculated by first computing the mean number of cells per slice for each subject, then averaging across all subjects.

#### Primary NG neuron isolation and culture

Rats were deeply anesthetized with 5% isoflurane and sacrificed via rapid decapitation. The left and right NG were dissected and placed in separate 1.5 mL Eppendorf tubes containing 1 mL Hanks Balanced Salt Solution (HBSS). Samples were immediately used for cell culture.

Dissected samples in HBSS were transported to a biosafety cabinet. HBSS was removed from samples. NG were physically dissociated with scissors and incubated for 40 minutes in Dulbecco’s Modified Eagle Medium (DMEM) and 1 mg/mL collagenase type IV. Solution was removed, and NG then incubated for 45 minutes in DMEM and 1 mg/mL trypsin. NG were plated onto laminin-coated glass coverslips within a 12-well plate and incubated for 1 hour. Neurons were fed with media containing DMEM with 10% heat-inactivated horse serum, 2 mM L-glutamine, 1% glucose, and 100 units/mL pen-strep. Samples were incubated at 37 °C overnight and imaged the next day.

#### Calcium imaging

Calcium imaging was performed on neurons using the ratiometric calcium indicator dye Fura-2-AM. Neurons were washed with 2.5 µg/mL Fura-2 in 2% BSA for 45 minutes at room temperature and then washed with extracellular buffer for 15 minutes. Extracellular buffer (7.4 pH, 320 Osm) contained 150 mM NaCl, 10 mM HEPES, 8 mM glucose, 5.6 mM KCl, 2 mM CaCl_2_, and 1 mM MgCl_2_. Coverslips were mounted onto an inverted fluorescence microscope and perfused with room temperature extracellular buffer at a rate of 6 mL/min. Cells were perfused with extracellular buffer for 1.5 minutes, A71623 for 2 minutes (100 nM in extracellular buffer), extracellular buffer for 2 minutes, KCl for 1 minute (50 mM in extracellular buffer), and extracellular buffer for 30 seconds. Fluorescence images were captured at 340 and 380 nm every 0.5 seconds.

#### Calcium imaging analysis

Cells were excluded from further analysis if there was a greater than 25% increase in the 340/380 nm ratio during the last 60 seconds of the initial extracellular buffer perfusion. The 340/380 nm ratio traces were then smoothed using a 15-point moving average filter prior to testing for drug responses. Cells that exhibited a greater than 25% increase in the 340/380 nm ratio during both A71623 and KCl treatment periods, compared to the 3.5 second pre-treatment windows, were counted as responders. Cells were considered non-responders to A71623 if they exhibited a greater than 25% increase in the 340/380 nm ratio during the KCl treatment period only. Cells that failed to respond to KCl treatment were excluded from analysis. For A71623-responsive neurons, a 3-point running average was computed for the ±1 sec period around the maximum raw 340/380 nm ratio during the A71623 treatment period. The maximum value of the 3-point smoothed data within this ±1 sec window was taken as the peak response. A chi-squared test was used to test whether there was a difference in the frequency of A71623-responsive neurons from left vs. right NG cultures.

### QUANTIFICATION AND STATISTICAL ANALYSIS

Data were preprocessed using Trailmaker™ (Parse Biosciences) to obtain cell-gene count matrices for each replicate group. Sequencing data were then analyzed with R (version 4.4.1) using Seurat (version 4.3.0). For the pseudobulk analysis, statistically significant left-right differential gene expression is reported for genes with at least a 1.5-fold difference in gene expression between the two sides, and a Bonferroni corrected p < 0.01. To test for lateralized cell counts for each identified cluster in the rat dataset, Bonferroni-corrected binomial tests were performed. For each cluster identified as having significant left or right-side biased counts (corrected p < 0.05), additional binomial tests were used to validate that this bias was observed in samples from both males and females. In the rat + mouse integrated dataset, we tested for lateralized expression of our *Chrna3* and *Cckar* co-expressing cluster of interest using a binomial test. Statistical significance is reported for p < 0.05.

For analysis of RNA-FISH data, paired t-tests were used to test for differential left-right expression of labeled neurons. Statistical significance is reported for p < 0.05. MATLAB was used to plot data and perform statistical tests.

In the calcium imaging experiments, a chi-squared test was used to test whether there was a difference in the frequency of A71623-responsive neurons from left vs. right NG cultures (GraphPad Prism). Peak amplitudes for left vs. right A71623-responsive neurons were compared using an unpaired t-test (MATLAB). Statistical significance is reported for p < 0.05.

Unless otherwise noted, summary data are presented as mean ± SEM. Additional statistical details, including precise p values and numerical n values, can be found in the text and figure legends.

---

**Figure S1.**
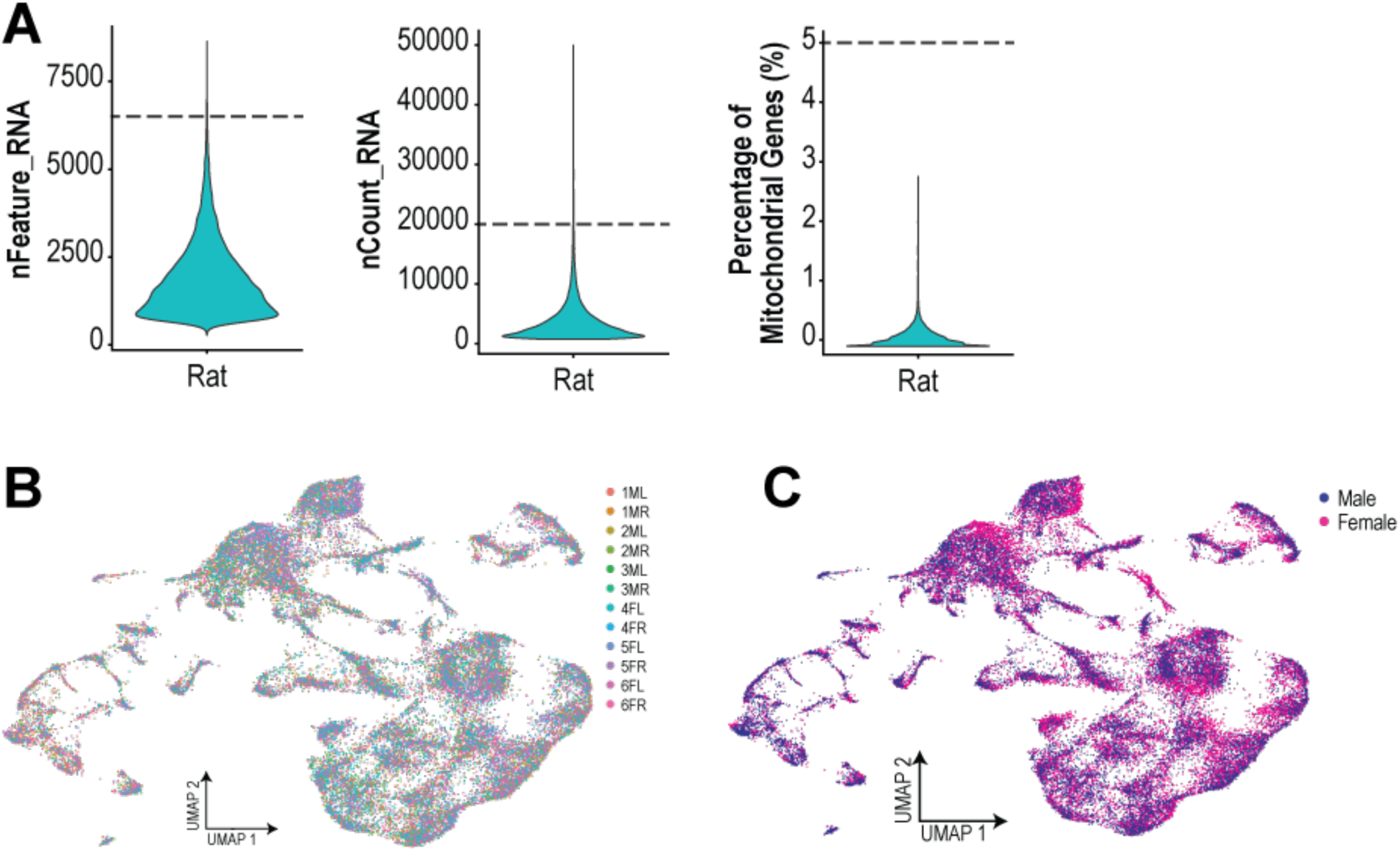
Quality control filtering and integration of replicate groups for rat dataset, Related to Figure 1. A) Nuclei with greater than 6,500 features (left), greater than 20,000 molecules (center), or greater than 5% mitochondrial genes (right) were removed prior to clustering. Before filtering, mean nFeature_RNA = 1857.668, mean nCount_RNA = 3804.594, and mean mitochondrial gene percentage = 0.179%. B) Samples from all 12 replicates (3 each from Male|Left (ML), Male|Right (MR), Female|Left (FL), and Female|Right (FR) treatment groups) were integrated prior to clustering. C) Samples from male and female rats exhibited overlapping UMAP profiles after sample integration.

**Figure S2.**
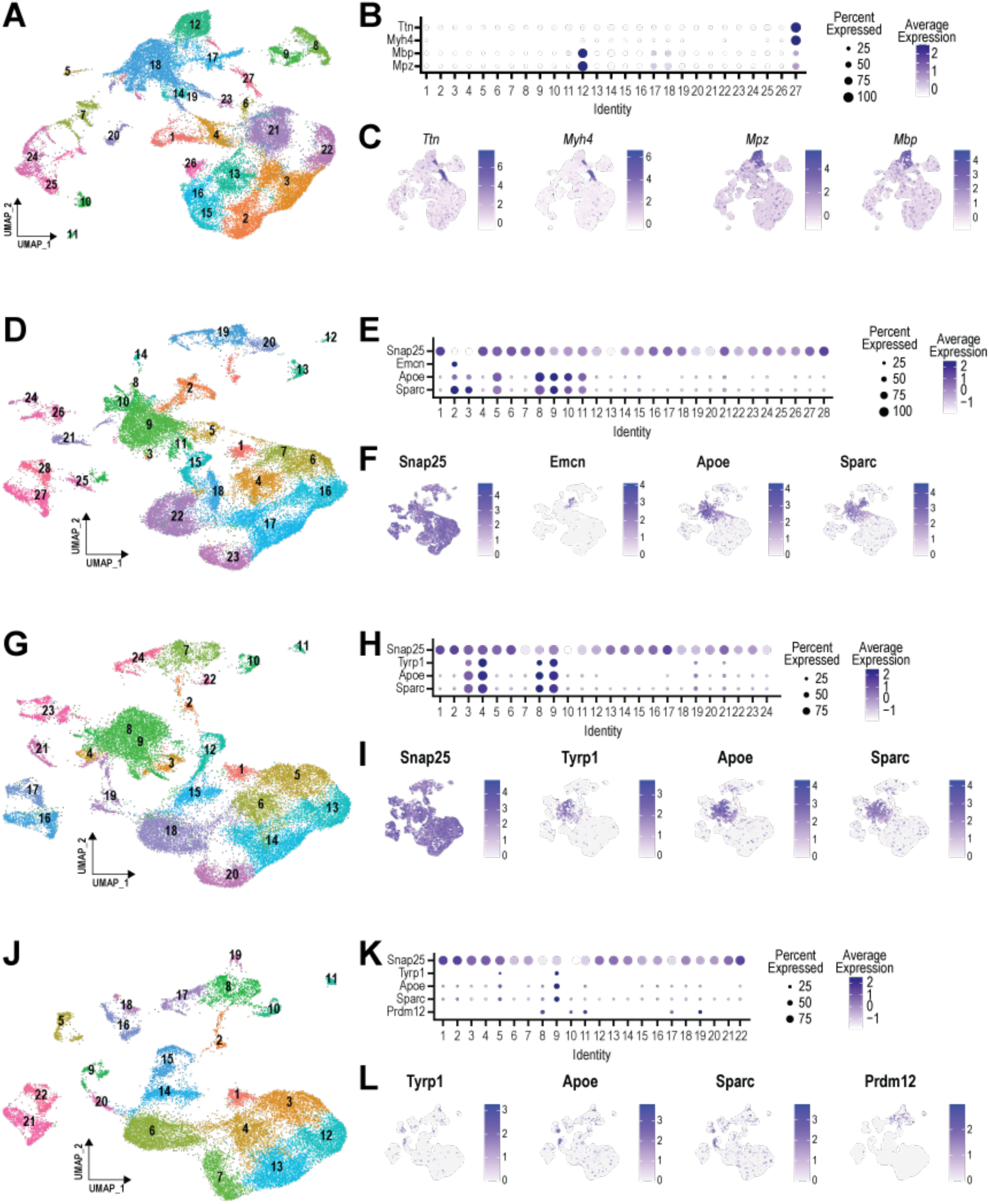
Filtering of clusters representing non-neuronal subtypes and jugular nodose neurons in rat data, Related to Figure 1. Four rounds of clustering and filtering were performed. A-C) In Round 1, two of 27 clusters were removed based on strong expression of marker genes associated with striated muscle (*Ttn*, *Myh4*; cluster 27) and myelin (*Mpz*, *Mbp*; cluster 12). A: Round 1 UMAP; B: dotplot showing expression of *Ttn*, *Myh4*, *Mpz*, and *Mbp* within each cluster; C: feature plots showing expression of targeted marker genes within filtered clusters. D-E) As in A-C for Round 2, in which 2 of 28 clusters were removed based on weak expression of the neuronal marker *Snap25* in conjunction with strong expression of the endothelial marker gene *Emcn* (cluster 2) and/or strong expression of the satellite glial marker genes *Apoe* and *Sparc* (clusters 2 & 3). G-I) As in A-C for Round 3 of cluster filtering, in which 4 of 24 clusters were removed based on strong expression of the melanocyte marker gene *Tyrp1* in conjunction with *Apoe* and *Sparc* (clusters 3, 4, 8, & 9). J-L) As in A-C for Round 4 of cluster filtering, in which 1 of 22 clusters was removed based on strong co-expression of *Tyrp*, *Apoe*, and *Sparc* (cluster 9), and 3 were removed based on strong expression of the jugular ganglia neuronal marker gene *Pdrm12*.

**Figure S3.**
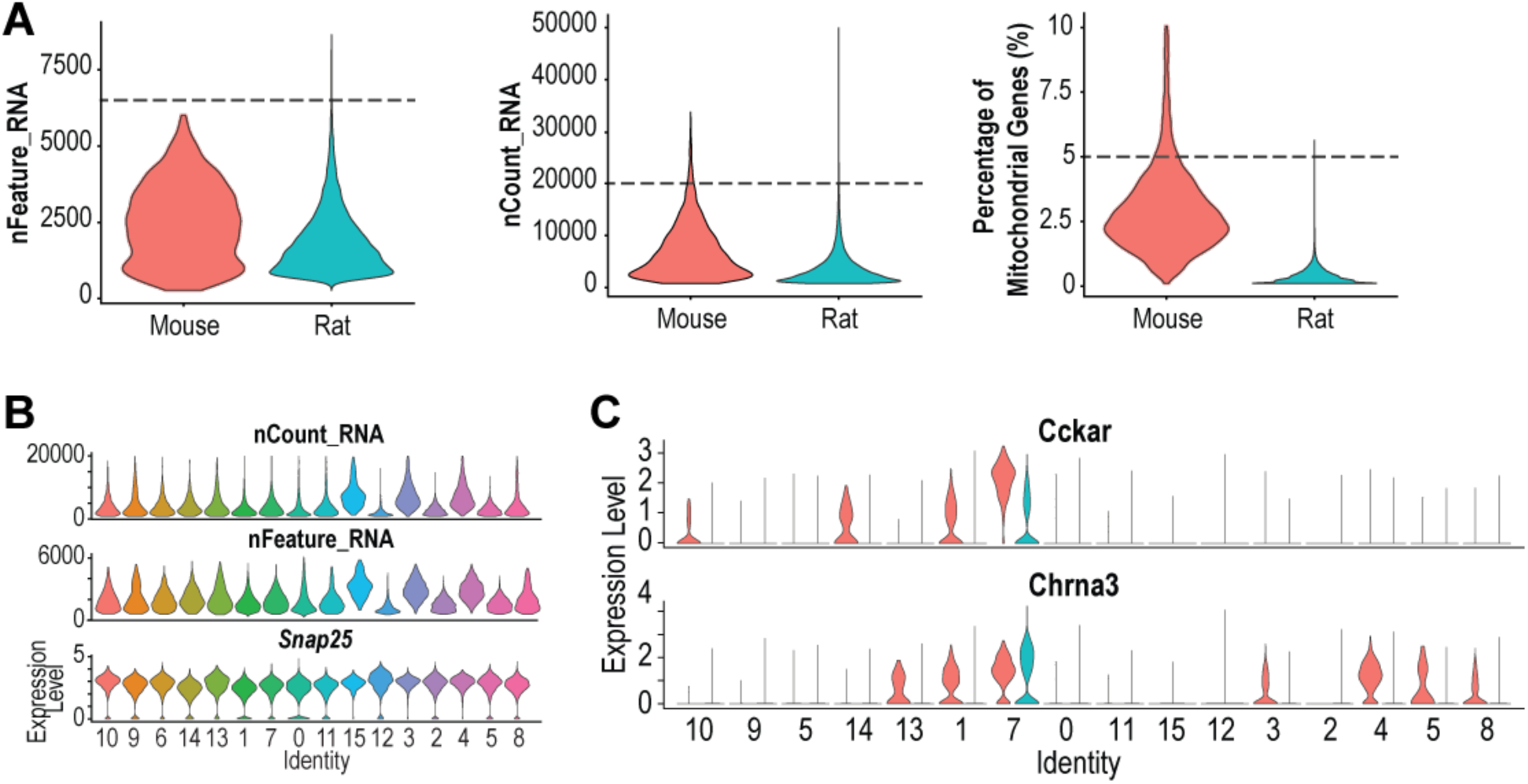
Quality control filtering of the rat + mouse integrated dataset, Related to Figure 6. A) For both datasets, nuclei with greater than 6,500 features (left), greater than 20,000 molecules (center), or greater than 5% mitochondrial genes (right) were removed prior to clustering. Dashed lines denote QC thresholds for each metric. Before filtering, mouse (red) mean nFeature_RNA = 2858.082, mean nCount_RNA = 7851.702, and mean percent mitochondrial genes = 2.922%. Rat data (blue) is as in Figure S1. B) After QC filtering and removal of non-neuronal cells and jugular ganglia neurons, remaining clusters all met quality control criteria and exhibited strong expression of the neuronal marker *Snap25*. C) Cluster 7 was the only cluster that exhibited expression of both *Cckar* and *Chrna3* in rats, and co-expression of these genes was likewise highest in mice.

**Figure S4.**
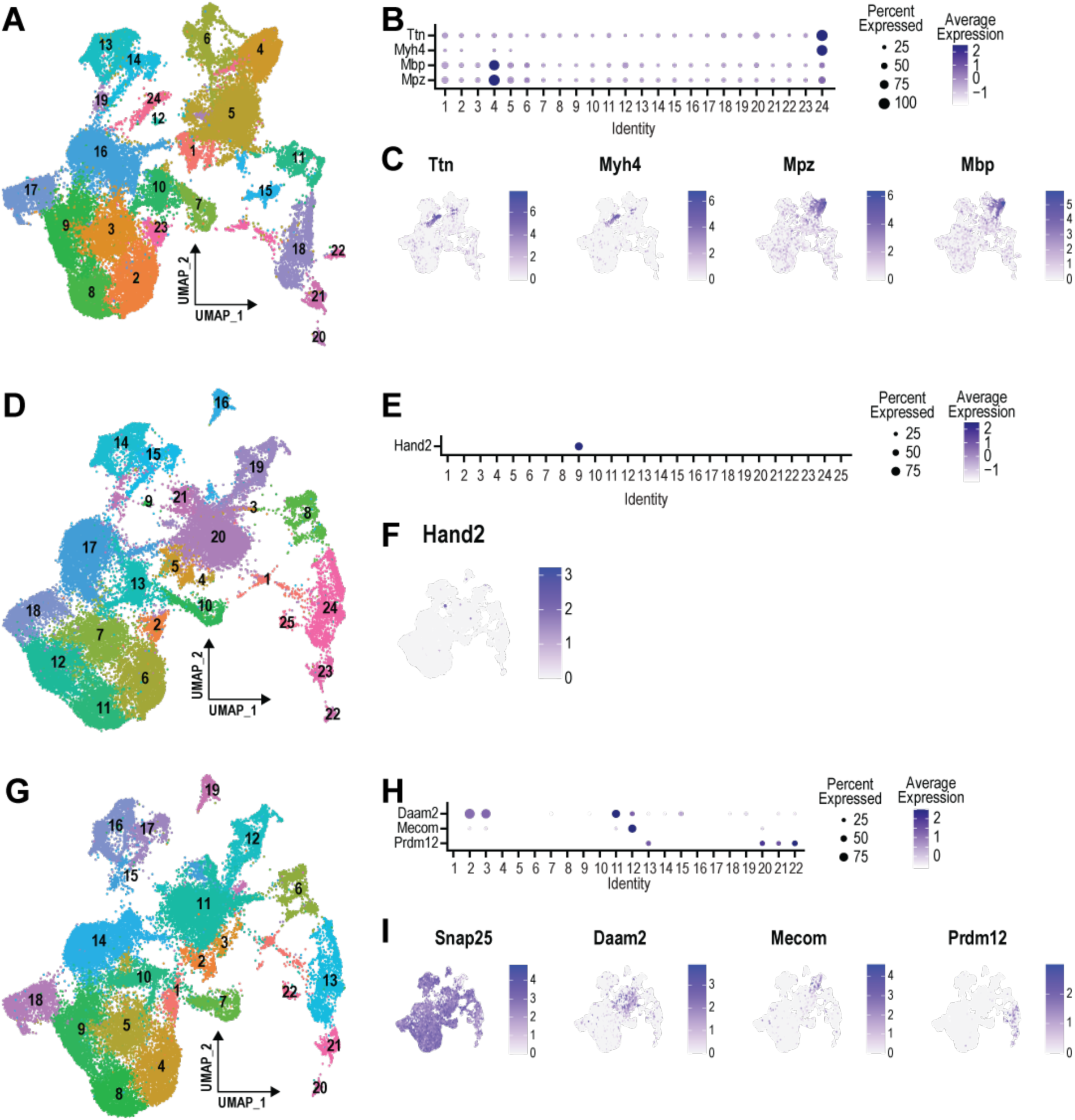
Filtering of clusters representing non-neuronal subtypes and jugular nodose neurons in the integrated rat + mouse dataset, Related to Figure 6. Three rounds of clustering and filtering were performed. A-C) In Round 1, two of 24 clusters were removed based on strong expression of marker genes associated with striated muscle (*Ttn*, *Myh4*; cluster 24) and myelin (*Mpz*, *Mbp*; cluster 4). A: Round 1 UMAP; B: dotplot showing expression of *Ttn*, *Myh4*, *Mpz*, and *Mbp* within each cluster; C: feature plots showing expression of targeted marker genes within filtered clusters. D-F) As in A-C for Round 2, in which 1 of 25 clusters was removed based on strong expression of the sympathetic neuron marker gene *Emcn* (cluster 9). G-I) As in A-C for Round 3 of cluster filtering, in which 1 of 22 clusters was removed based on strong expression of the endothelial marker gene *Mecom*, 1 cluster was removed based on strong expression of the oligodendrocyte marker gene *Daam2*, and 4 clusters were removed based on expression of the jugular ganglia neuronal marker gene *Prdm12* (clusters 13, 20, 21, and 22).

## Notes

https://www.ncbi.nlm.nih.gov/geo/query/acc.cgi?acc=GSE280062

### Competing Interest Statement

The authors have declared no competing interest.

### Summary of Updates

We have re-analyzed our sequencing data, we have added new in situ hybridization and calcium imaging experiments to demonstrate functional asymmetry in expression and signaling within our identified cell population, and we have completely rewritten the manuscript to present these new results and provide additional clarity. Our fundamental finding remains unaltered. As before, we demonstrate that there is a unique population of Cckar-expressing vagal neurons that is preferentially found in right nodose ganglia over left. Consistent with these results, we now show that CCK receptor agonism activates a significantly greater percentage of right nodose ganglia neurons than left, providing novel evidence of functional left-right asymmetry in peptidergic vagal signaling. We have added an author, added Figure 7 and caption, and added Supplemental Figures. All existing text and figures were revised.

